# Full-Term and Preterm Differences in Human Brain Functional Connectivity Development Across Birth

**DOI:** 10.64898/2026.10.29.762902

**Authors:** Lanxin Ji, Devyn Cotter, Sarah Banker, Christopher J. Trentacosta, Moriah E. Thomason

## Abstract

We possess a limited understanding of how preterm and full-term brains develop during the earliest stages of life, from the fetal to neonatal period, and how they respond to the birth transition. Here, we characterize brain resting-state functional connectivity (RSFC) using over 1000 longitudinal fetal and infant functional MRI scans from 21 to 52 post-conception weeks across birth. By mapping the developmental trajectory of RSFC across this window, we unveil marked divergences in developmental patterns between preterm- and term-born brains at both the global and network levels. When aligned by birth, the full-term group exhibit generally higher RSFC than the preterm group. For selected between-network connections with a significant prematurity effect, full-term cases demonstrate an overall increase in RSFC across age, whereas preterm cases show minimal change or even a decrease in connectivity of these systems. In contrast, when aligned by postconceptional age, the preterm group shows an accelerated increase in RSFC between 30 and 40 weeks. Across comparisons, mode of delivery and sex showed minimal influence on these trajectories. These findings offer novel insight into how prematurity and early exposure to the extrauterine environment alter the developmental trajectories of large-scale neural networks, advancing our understanding of both typical and preterm brain development.

## Introduction

Preterm birth is a significant public health concern, affecting approximately 10% of live births worldwide ^1^. The implications of being born early are profound, extending beyond the neonatal period throughout the life cycle. Preterm infants are at higher risk of a range of neurobehavioral and psychiatric conditions, including the inattentive subtype of attention-deficit/hyperactivity disorder (ADHD), depression, anxiety, autism spectrum disorder (ASD), avoidant personality disorder, and antisocial personality disorder ^2–4^. Children who were born preterm also show higher rates of language, cognitive, sensory, and motor impairments, which can manifest as poor academic performance or behavioral problems in school ^4–7^, though some differences may diminish with age ^8^.

The last trimester of pregnancy represents a critical period in fetal brain development. Brain growth peaks and nerve cells form, multiply, and migrate to create a vast network of connections ^9,10^. In the context of prematurity, it is possible that a newborn being introduced to the ex-uterine environment prior to the typical peak in brain maturation that occurs around term, could lead to a markedly different developmental strategy. Studying how prematurity interrupts this essential transition from the fetal to the neonatal period is critical for understanding the mechanisms underlying adverse outcomes.

Studies using resting-state functional Magnetic Resonance Imaging (fMRI) have shown altered brain functional patterns in preterm infants compared to term controls. For example, prominent reductions in resting-state functional connectivity (RSFC) were noted in preterm infants between the thalamus and cortical regions including the sensorimotor, prefrontal, insular, and anterior cingulate cortices ^11^, as well as between the amygdala and subcortical or brainstem structures, including the thalamus, hypothalamus, brainstem, and insula ^12^. Altered RSFC in subcortical and cortical networks has been linked to cognitive, motor, emotional, and behavioral outcomes in early childhood ^13–15^. Large-scale connectome analyses reveal striking deficiencies in within-network connectivity across canonical resting-state networks, with earlier gestational age at birth associated with weakened RSFC ^16^. These impairments are characterized by a 23–41% reduction in core network strength, including motor, sensory, visual, auditory, frontal, and temporal networks, accompanied by paradoxical increases in bilateral superior parietal lobule connectivity within the lateral motor network ^16^. Despite the reductions in network strength, preterm infants exhibit preserved small-world topology and modular organization^17,18^. Infants born very preterm show reduced segregation and integration at term-equivalent age compared to term controls, but their RSFC within the occipital/visual module is increased ^19^. In brain analysis, segregation refers to the specialized functioning of distinct brain regions, whereas integration describes the coordinated activity across different brain networks. Other work on dynamic functional connectivity analyses indicates that the neonatal brain expresses multiple connectivity states whose temporal properties are altered in preterm infants, and that these alterations are linked to atypical social, sensory, and repetitive behaviors at 18 months ^20–22^.

It is possible that prematurity-related diruptions in brain connectivty arises even earlier, prior to birth. A rare fetal cohort study demonstrated that fetuses destined for preterm delivery show decreased segregation between language areas compared to those delivered at term ^23^, providing the first evidence that prematurity-related brain alterations begin in utero, before postnatal environmental factors come into play. However, another study showed that when premature infants are compared to conceptional-age-matched fetuses, the relationship reverses: they appear more mature^24^. The challenge that arises when comparing fetal and newborn brains is that it is difficult to disambiguate the consequences of exogenous environmental input on brain maturation. Thus, available cross-sectional studies have yet to isolate inflection points or individual trajectories that may be vitally important for understanding large-scale circuit organization and later outcomes in infants born preterm.

Developing a model capable of capturing this dynamic requires a large, densely sampled longitudinal dataset that spans the prenatal-to-postnatal transition. To address this need, we mapped trajectories of functional circuit development in over 1,000 fMRI scans from term- and preterm-born fetuses and neonatess, combining the public data from the Developing Human Connectome Project (dHCP) and our own longitudinal cohort, the Perinatal Imaging of Neural Connectivity (PINC) project. To our knowledge, this combined dataset represents the largest perinatal neuroimaging sample in the world. We expect to differentiate preterm from full-term developmental trajectories based on altered timing of events or changes in trajectory shape. To disentangle the mixed effects of continuous development from conception and potential discontinuities introduced by birth, we examine brain functional connectivity using two complementary temporal frameworks: post-conception gestational age (GA), indexing the intrinsic biological maturation linked to gestational timing since conception, and birth-centered age, which centers birth at zero to highlight neural reorganization associated with the birth event and subsequent ex-utero exposure. Comparing developmental trajectories in both frameworks allowed us to distinguish whether differences observed in preterm cases arose from altered maturational pacing or from adaptations to the environmental transition at birth. We also examine the mode of delivery as a potential contributor to developmental differences. Key questions addressed by this work include: (1) how preterm neonates diverge from their full-term counterparts in brain development cross birth, (2) whether differences in brain developmental trajectories between full-term and preterm cases vary across brain regions, reflecting region-specific patterns of development and adaptation, and (3) whether mode of delivery is reflected in neural maturational differences.

## Results

### Characteristics of the final sample

The final sample comprised n = 1,031 fMRI scans, including 244 preterm and 787 full-term cases, with detailed demographic and imaging quality parameters summarized in **Table 1**. Birth before 37 weeks of gestation is defined as preterm. Imaging quality was quantified by the root-mean-square of voxel-wise differentiated signal (DVARS, ^25^). As significant between-group differences were observed in the imaging quality metric, a sensitivity analysis using a subset of data matched for imaging quality is provided in the Supplementary Materials **Table S1** and **Figure S1**.

**Table 1.**
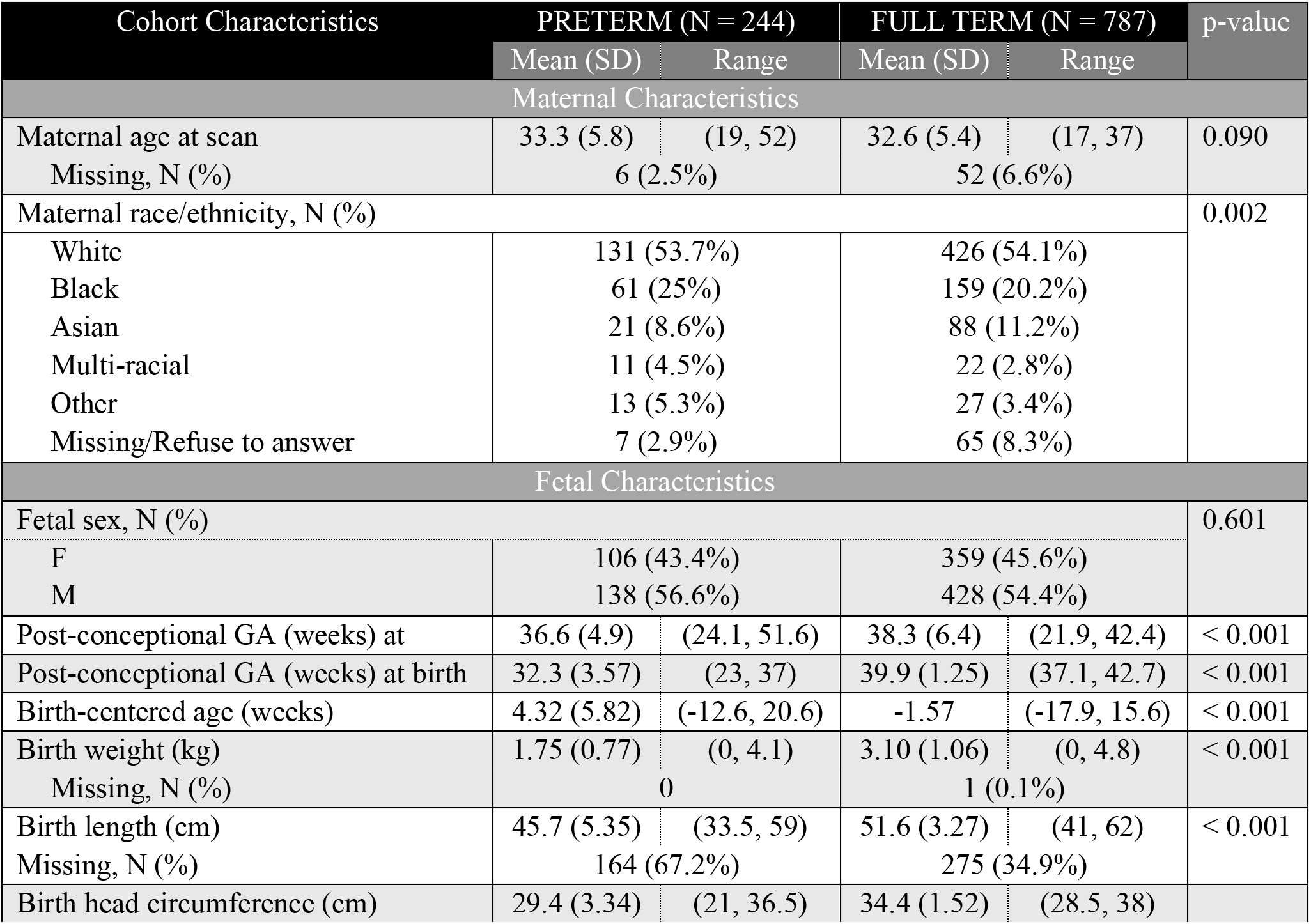
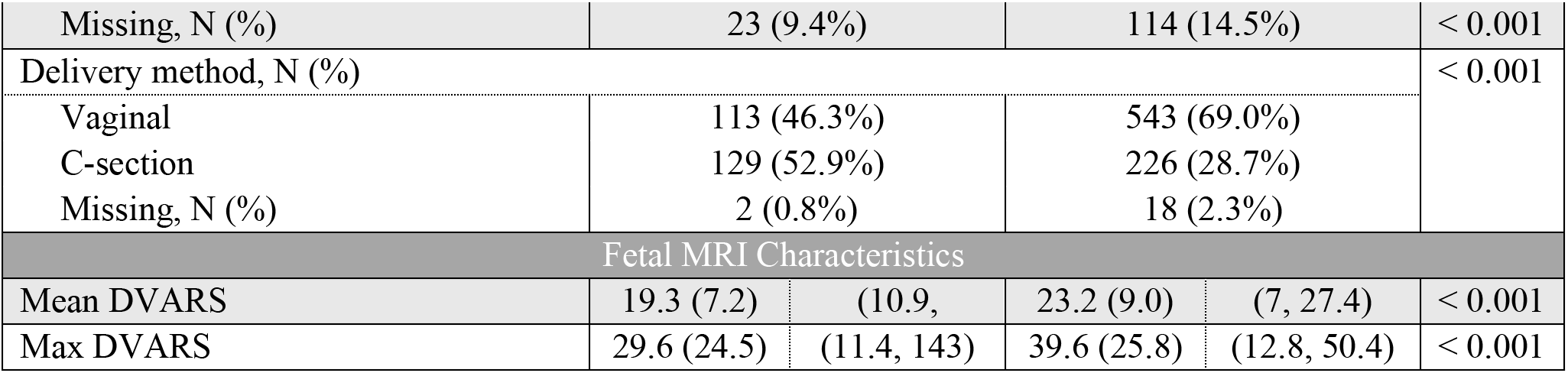
Group comparisons across demographic factors, birth outcomes, and MRI data quality.

### Full-term show higher RSFC than preterm cases across the birth transtion period

Preprocessed fetal and infant fMRI data were used to generate a data-driven, group-balanced functional atlas comprising 200 functional parcels within the gray matter. For each region of interest (ROI), the voxel-wise time series were averaged to produce a single representative time series, which was then used to construct a 200 × 200 connectivity matrix for each scan. Age-related RSFC was then isolated using regression models across all subjects, controlling for group-related variance in age at scan.

We first aligned the data by birth-centered age. Among the 19,900 possible ROI-to-ROI RSFC pairs, 22.6% showed a significant increase in RSFC with birth-centered age, while 28.2% showed a decrease with age. Developmental trajectories of RSFC were examined at the global level by averaging RSFC values showing significant positive or negative associations with age to generate growth plots. To quantify periods of significant developmental change and estimate developmental rates, the first derivative (local slope) of each GAMM-fited curve was calculated. As seen in **Figure 1d-e**, Full-term cases displayed a sharp postnatal change at birth for both positive and negative age-related RSFC, marking birth as an inflection point in functional connectivity development. Preterm infants, however, showed a more gradual, near-linear increase or decrease. That is, the acceleration in connectivity seen in term infants leading up to and following birth was not evident in those born preterm. In the regression model combining all data (RSFC ∼*s*(Age) + Preterm + Preterm × Age + (1∣Subject)), both preterm status and the Preterm × Age interaction were significantly associated with functional connectivity for connections exhibiting either positive or negative relationships with birth-centered age.

**Figure 1.**
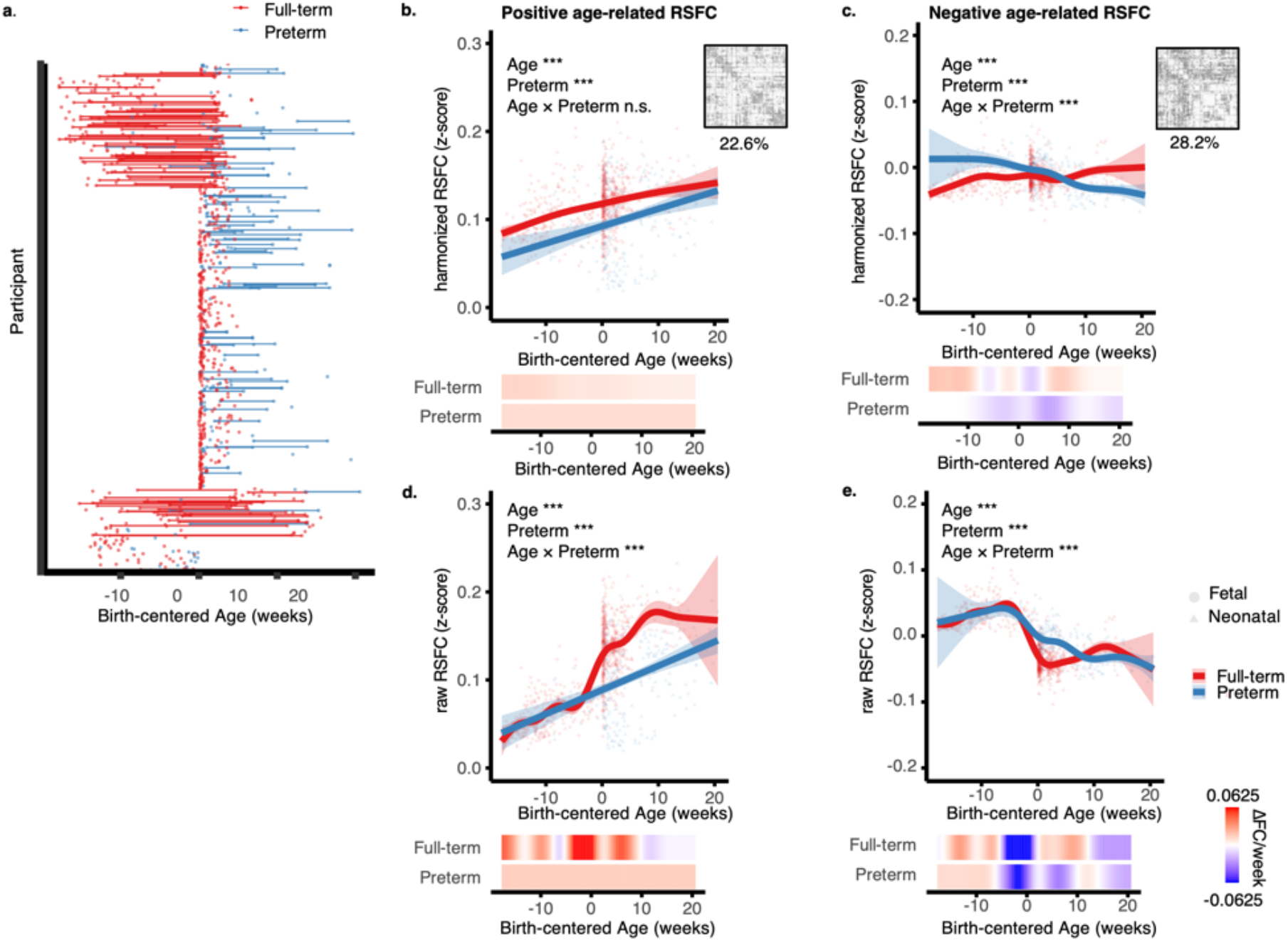
Longitudinal trajectories of functional connectivity development across birth as a function of birth-centered age. (a) Lollipop plot showing participant timelines. Each row represents one participant; points indicate scan time. Lines and points are color-coded by group (full-term: red, preterm: blue). (b–c) Developmental trajectories and corresponding rates of change for average GA-related RSFC following harmonization to account for batch effects (positive age-relations in b, negative in c) in full-term and preterm cases. Curves and data points are shown in red and blue for full-term and preterm groups, respectively. Local rates of RSFC change are displayed below each panel on a blue-to-red scale. (d–e) Developmental trajectories and rates of change for GA-related raw RSFC z-scores without harmonization (positive age-relations in e, negative in f) in full-term and preterm infants. Conventions for curves, confidence intervals, symbols, and local rate maps follow those in b–c. Shaded bands denote 95% confidence intervals. Fetal and infant scans are marked by triangles and circles, respectively. Signif. codes: 0 ‘***’ 0.001 ‘**’ 0.01 ‘*’ 0.05 ‘.’ 0.1 ‘n.s’ 1.

A critical challenge was distinguishing biological changes across the fetal-to-infant transition from differences introduced by imaging acquisition and processing protocols. To account for dataset differences, Longitudinal ComBat^26^ was applied to harmonize age-related RSFC across acquisition sequences. Trajectories were modeled using generalized additive mixed-effect models (GAMMs), following the equation RSFC ∼ 1 + s(GA) + (1|Subject), separately for full-term and preterm infants. Post-harmonization, the significant difference between preterm and full-term groups effect was maintained, although the burst increase at birth disappeared, which could be related to over-correction of MRI condition differences, as MRI collection setting (in utero vs. ex utero) is fully aligned with the biological effect of birth that we aim to study. Notably, we again observed higher RSFC values in the full-term group than in the preterm group (**Figure 1b-c**).

### Indicators of gestationally programmed development reveal accelerated increase during the 3^rd^ trimester in preterm cases

Functional connectivity analysis and regression modeling of post-conceptional GA followed the same procedures as described for birth-centered age. When aligned by post-conceptual GA, 27.8% of ROI-to-ROI RSFC pairs showed a significant increase with post-conceptional GA, while 16.9% showed a decrease with GA. RSFC followed a near-linear trend in positive age-related RSFC and a nonlinear pattern in negative age-related RSFC across gestation and early postnatal life (**Figure 2d-e**). Post-harmonization, preterm cases exhibited an accelerated increase in RSFC between 30 and 40 post-conceptional weeks (**Figure 2b-c**), the window during which preterm deliveries typically occur, whereas full-term cases showed a steady increase across the entire transition period. Notably, preterm birth had a significant effect on RSFC changes after harmonization, but not in the raw RSFC data.

**Figure 2.**
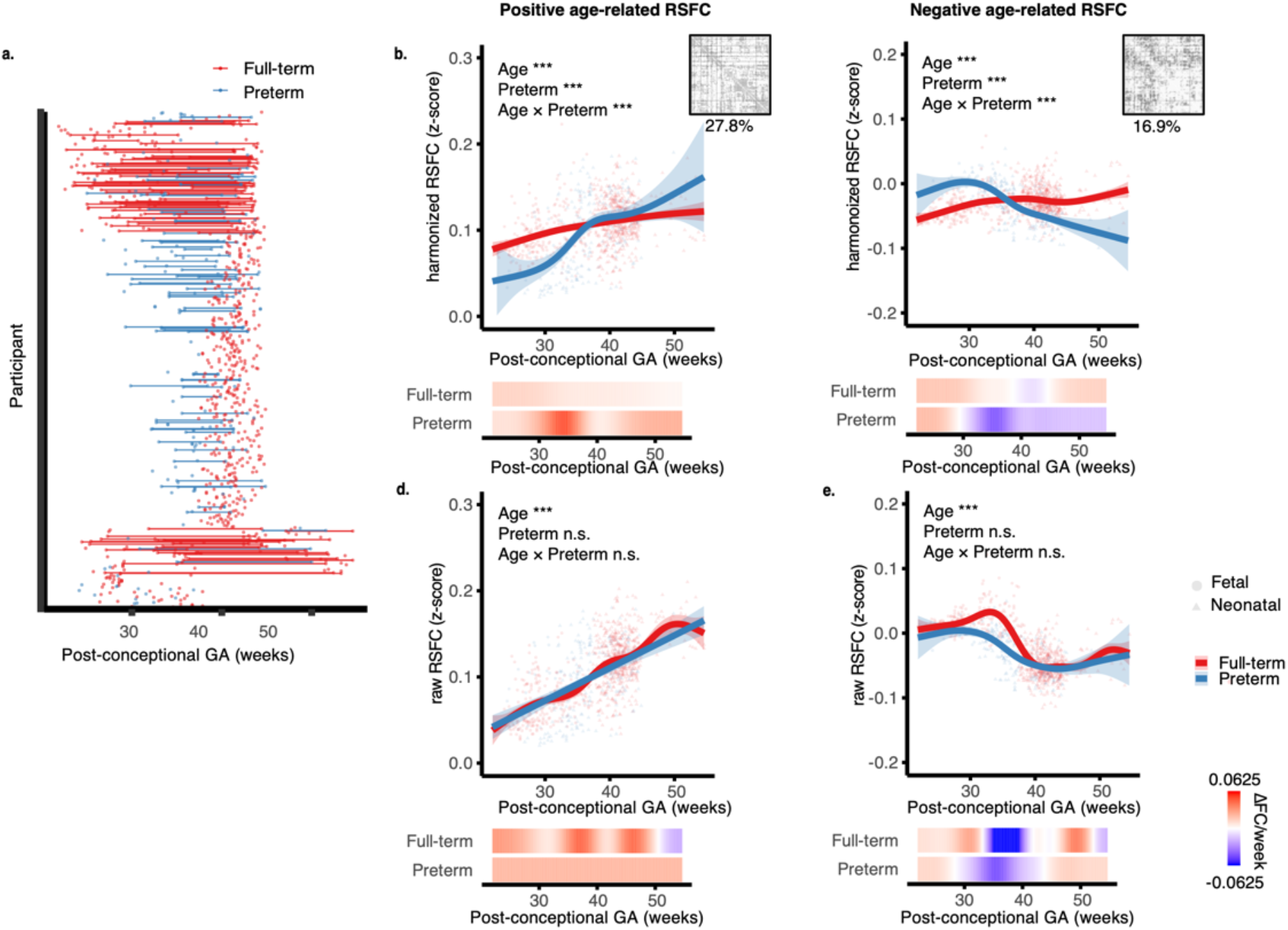
Longitudinal trajectories of functional connectivity development across birth as a function of post-conceptual GA. (a) Lollipop plot showing participant timelines. Each row represents one participant; points indicate scan time. Lines and points are color-coded by group (full-term: red, preterm: blue). (b–c) Developmental trajectories and corresponding rates of change for average GA-related RSFC following harmonization to account for batch effects (positive age-relations in b, negative in c) in full-term and preterm cases. Curves and data points are shown in red and blue for full-term and preterm groups, respectively. Local rates of RSFC change are displayed below each panel on a blue-to-red scale. (d–e) Developmental trajectories and rates of change for GA-related raw RSFC z-scores without harmonization (positive age-relations in e, negative in f) in full-term and preterm infants. Conventions for curves, confidence intervals, symbols, and local rate maps follow those in b–c. Shaded bands denote 95% confidence intervals. Fetal and infant scans are marked by triangles and circles, respectively. Signif. codes: 0 ‘***’ 0.001 ‘**’ 0.01 ‘*’ 0.05 ‘.’ 0.1 ‘n.s’ 1

### Network-specific differences in brain related to birth

Using a hierarchical clustering algorithm within the CONN toolbox ^27^, 8 functional networks emerged based on the similarity of their RSFC patterns (**Figure 3a**). At the perinatal stage, the networks contained bilaterally distributed regions and some resembled primitive, adult-like functional networks. The identified networks included the Supplementary Motor Area (SMA), Cingulo-Parietal Network (CPN), Frontal-Parietal Network (FPN), Sensorimotor, Bilateral Insular (INS), Auditory-Temporal Network (ATN), Visuomotor Integration and Coordination Network (VIC), and Default Mode Network (DMN).

**Figure 3.**
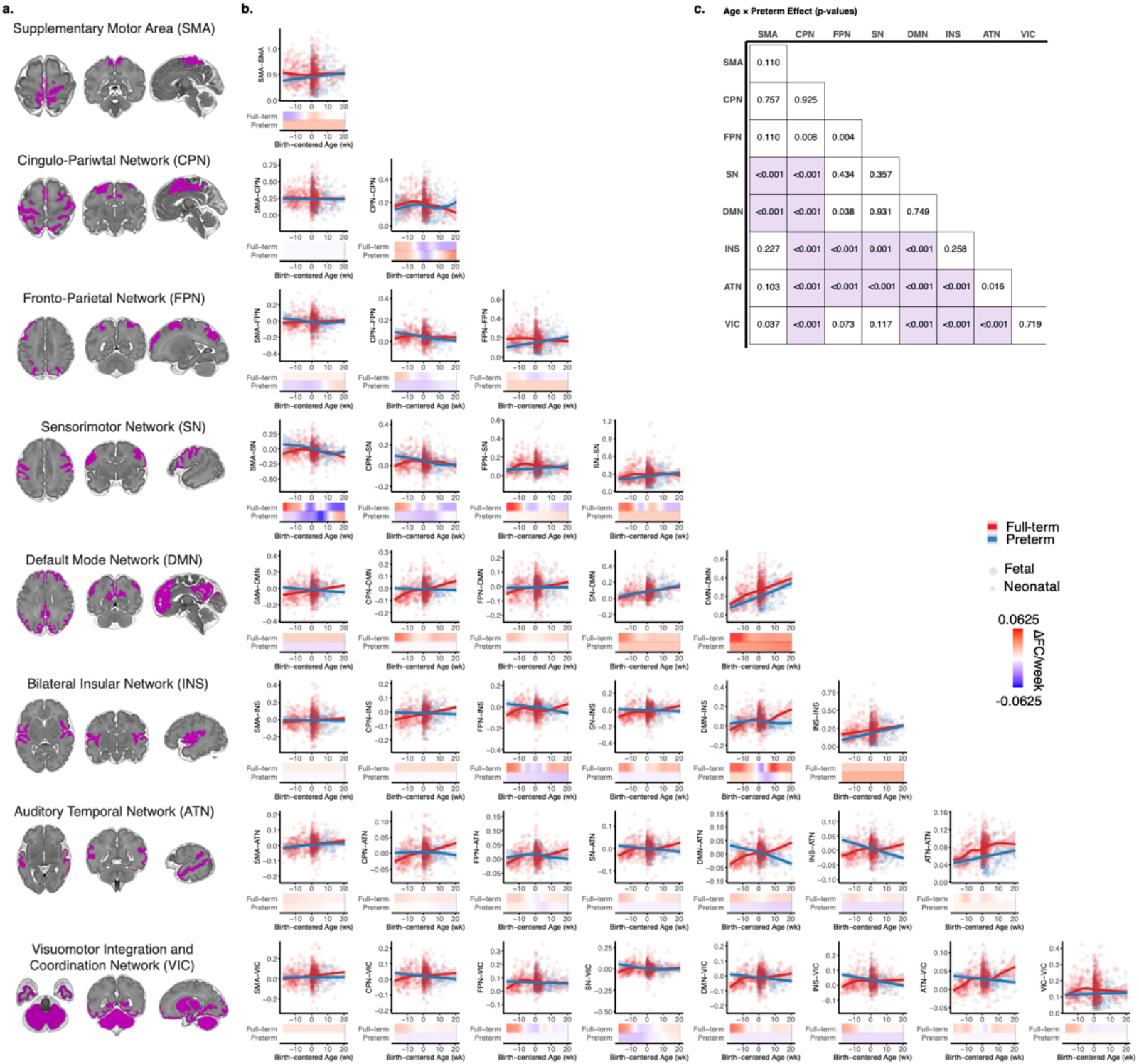
Differences in brain network RSFC between preterm and full-term infants in relation to birth-centered age. (a) Functional networks included Supplementary Motor Area (SMA), Cingulo-parietal (CPN), Fronto-parietal (FPN), Sensorimotor, Bilateral Insular (INS), Auditory Temporal (ATN), Visuomotor Integration (VIS), and Default Mode Network (DMN). (b) Developmental trajectories and corresponding rates of change for age-related RSFC in preterm and full-term groups. Curves and data points are colored red (full-term) and blue (preterm), with shaded regions indicating 95% confidence intervals. Local rates of change (first derivatives) are shown below each trajectory, color-mapped from blue (negative) to red (positive). Shaded bands denote 95% confidence intervals. Fetal and infant scans are marked by triangles and circles, respectively. (c) P-values for the Preterm × Age interaction effect on network-to-network RSFC. Connections showing a significant interaction (p < 0.001, Bonferroni-corrected) are highlighted in purple.

Network-level RSFC was derived by averaging ROI-to-ROI connections that showed significant positive or negative associations with age. The network-level RSFC was calculated by averaging the ROI-by-ROI RSFC values that showed significant positive or negative associations with age. Here, positive and negative effects are averaged to depict dominant effects (e.g., positive or negative trending, overall) for each network.

The effect of prematurity was assessed using a GAMM: RSFC ∼*s*(Age) + Preterm + Preterm × Age + (1∣Subject). Using a birth-centered age framework, the developmental trajectories for each network-by-network pair in the preterm and full-term groups are shown separately in **Figure 3b**, along with the p-values for the Preterm × Age term in **Figure 3c**. Here, we observe that network-level RSFC differences between preterm and full-term cases occur predominantly in between-network connectivity, with the SN, DMN, INS, ATN, and VIC involved in these differences. A consistent pattern among the network-by-network connections showing a significant preterm effect is that the full-term group exhibit an overall increase in RSFC across the birth transition period, whereas the preterm show minimal change or even a decrease in RSFC in these systems over age.

**Figure 3** presents results based on post-harmonization data; trajectories derived from the pre-harmonized raw RSFC z-scores are provided in **Supplementary Figure S2**.

### Network-specific differences in maturational timing across post-conceptional gestational age

We repeated the above analyses using postconceptional GA age as the developmental index. As shown in **Figure 4**, prematurity was associated with a greater number of altered connections. Notably, all within-network RSFC measures, with the exception SN, exhibited a significant Preterm × GA interaction. Within-network RSFC in the SMA, CPN, FPN, ATN, and VIS networks showed developmental trajectories that closely paralleled the global pattern in **Figure 2b**, with preterm infants exhibiting an accelerated increase between 30 and 40 weeks’ GA. Although all networks contributed to preterm–full-term differences, the default mode network (DMN) was distinctive: all DMN-related connections, except DMN–SN RSFC, showed significant prematurity effects. In parallel, for the between-network connections showing preterm vs. full-term differences, the two groups tend to exhibit opposite developmental trajectories: the full-term group show increasing RSFC, whereas perterm show decreasing RSFC with post-conception GA.

**Figure 4.**
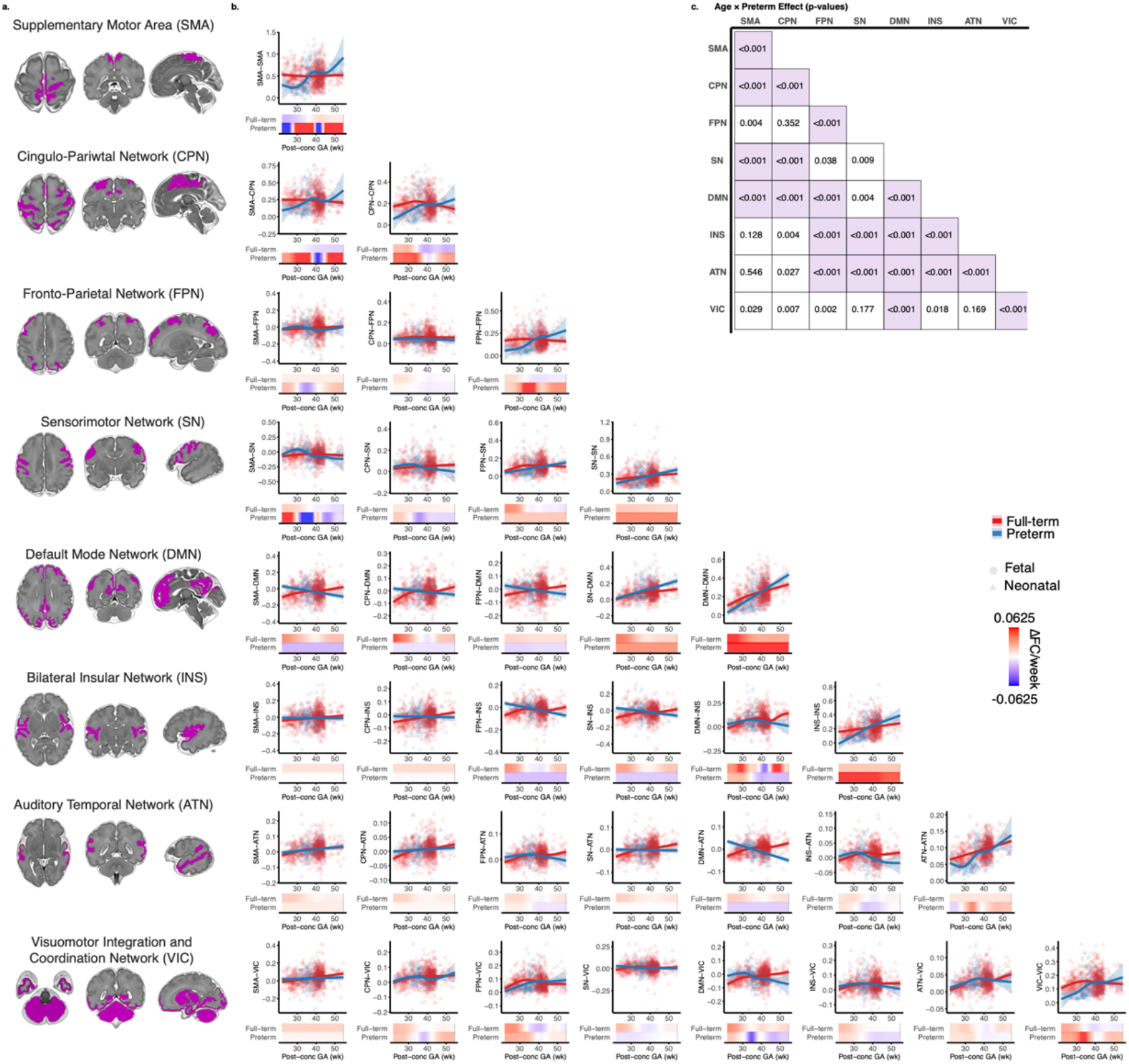
Differences in brain network RSFC between preterm and full-term infants in relation to post-conceptual gestational age (GA). (a) Functional networks. (b) Developmental trajectories and corresponding rates of change for GA-related RSFC in preterm and full-term infants. Curves and data points are colored red (full-term) and blue (preterm). Local rates of change (first derivatives) are shown below each trajectory, color-mapped from blue (negative) to red (positive). Shaded bands denote 95% confidence intervals. Fetal and infant scans are marked by triangles and circles, respectively. (c) P-values for the Preterm × Age interaction effect on network-to-network RSFC. Connections showing a significant interaction (p < 0.001, Bonferroni-corrected) are highlighted in purple.

Corresponding trajectories derived from pre-harmonized RSFC z-scores as a function of postconceptional GA are presented in **Supplementary Figure S3**.

### Effect of the Delivery Method

We further examined the potential effect of delivery method on brain RSFC changes surrounding birth. This analysis was performed only in the time framework of birth-centered age, as we do not expect the delivery method to relate to prenatal gestational programming. We applied a GAMM (RSFC ∼ s(Age) + Delivery + Delivery × Age + (1|Subject)) and conducted analyses separately in full-term and preterm infants, given that the influence of delivery mode may differ across these conditions. Overall, there were no major differences in the shape of RSFC developmental trajectories between those delivered vaginally and those by caesarean section (c-section)(Figure 5).

**Figure 5.**
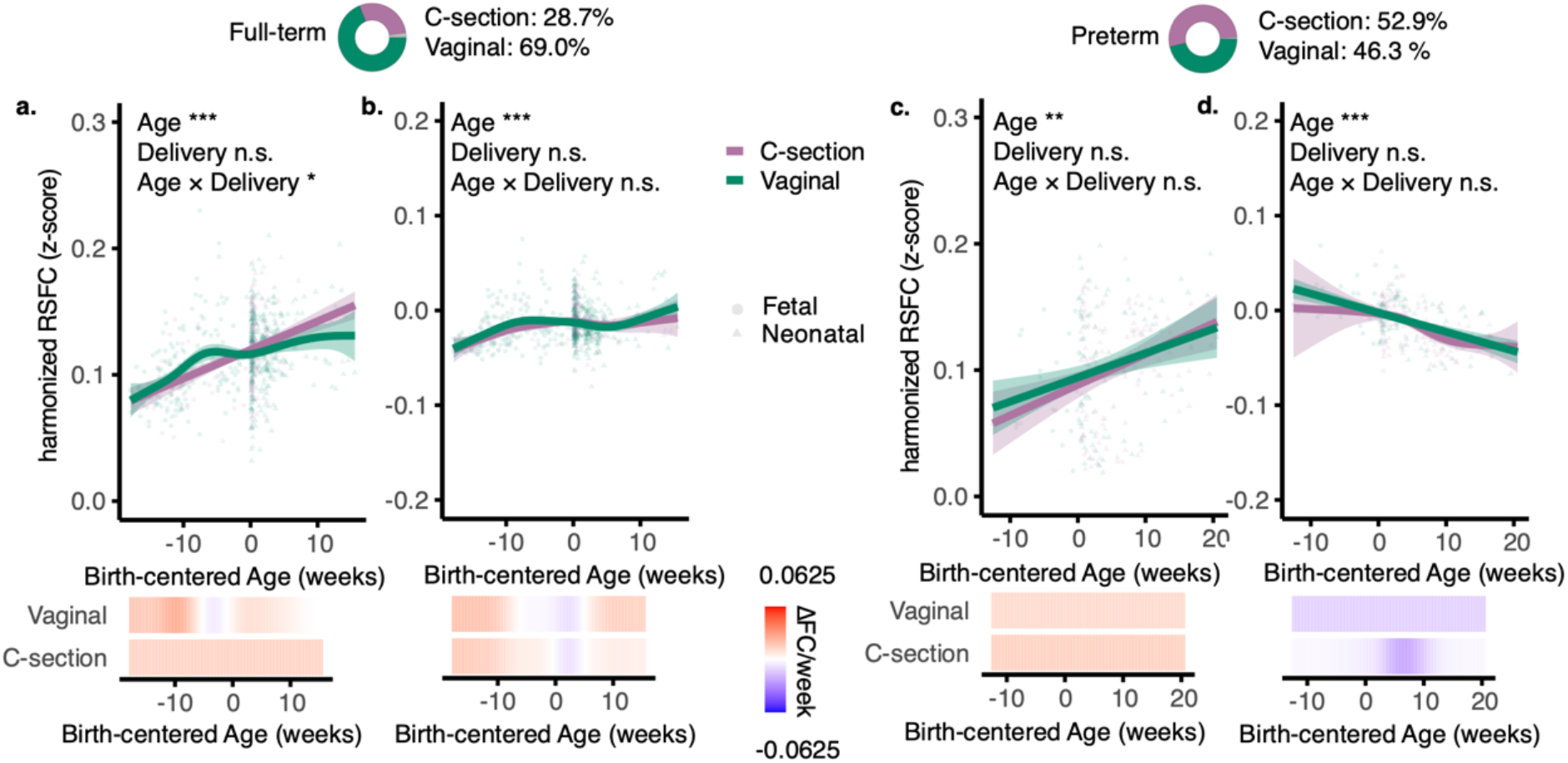
Developmental trajectories and rates of change in RSFC by delivery mode. (a) Full-term cases; (b) Preterm cases. Fitted curves and data points for infants with vaginal or caesarean section (c-section) delivery are shown in purple and green, respectively. The local rate of RSFC change is shown below the plot, color-coded from blue (negative) to red (positive). Shaded bands denote 95% confidence intervals. Fetal and infant scans are marked by triangles and circles, respectively.

A modest difference emerged in positive age-related RSFC changes across birth among full-term cases, with significant *Delivery × Age* interaction (*t* = –2.23, *p* = 0.029). Specifically, infants delivered by c-section showed a slightly greater postnatal, but less prenatal increase in RSFC compared with those delivered vaginally. In contrast, no significant effect of delivery method was observed in the preterm group. At the network level, no specific network-to-network RSFC pairs showed significant differences (after FDR correction) as a function of delivery method, either in the full-term or preterm group.

### Potential sex differences in brain functional development

We also assessed the potential influence of sex on RSFC changes across the birth transition. Using a GAMM framework (RSFC ∼ s(Age) + Sex + Sex × Age + (1|Subject)), we analyzed the full dataset as well as full-term and preterm groups separately. Across all analyses, we found no significant sex differences in RSFC, whether age was expressed as postconceptional GA or as birth-centered age. Coefficient test statistics are reported in **Table 2**. Likewise, no significant sex-by-age interactions were detected at the network level after FDR correction.

**Table 2.** Regression Coefficient Test Statistics for the Sex Effect.

| Model tests statistics | Postive age-related changes |  | Negative age-related changes |  |
| --- | --- | --- | --- | --- |
|  | t value | p value | t value | p value |
| All subjects |  |  |  |  |
| Post-conceptional GA at Scan | 8.849 | < 0.001 *** | 0.754 | 0.451 |
| Sex (Females >Males) | -1.101 | 0.271 | 0.315 | 0.753 |
| GA:Sex | 1.037 | 0.300 | -0.470 | 0.638 |
| Birth-centered Age at Scan | 4.404 | < 0.001 *** | 1.267 | 0.206 |
| Sex (Females >Males) | -0.006 | 0.995 | -1.465 | 0.143 |
| GA:Sex | 1.598 | 0.110 | -0.151 | 0.880 |
| Full-term |  |  |  |  |
| Post-conceptual GA at Scan | 6.304 | < 0.001 *** | 5.972 | < 0.001 *** |
| Sex (Females >Males) | -1.582 | 0.114 | 0.703 | 0.483 |
| GA:Sex | 1.525 | 0.128 | -0.978 | 0.328 |
| Birth-centered Age at Scan | 6.957 | < 0.001 *** | -5.040 | < 0.001 *** |
| Sex (Females >Males) | 0.404 | 0.686 | -1.899 | 0.058 |
| GA:Sex | 1.730 | 0.084 | -0.537 | 0.591 |
| Preterm |  |  |  |  |
| Post-conceptual GA at Scan | 7.184 | < 0.001 *** | -9.968 | < 0.001 *** |
| Sex (Females >Males) | -0.158 | 0.875 | 0.351 | 0.726 |
| GA:Sex | 0.113 | 0.910 | -0.286 | 0.775 |
| Birth-centered Age at Scan | 2.503 | 0.013 * | -6.528 | < 0.001 *** |
| Sex (Females >Males) | -1.004 | 0.316 | 0.857 | 0.392 |
| GA:Sex | 1.065 | 0.288 | -0.856 | 0.393 |

Signif. codes: 0 '\*\*\*' 0.001 '\*\*' 0.01 '\*' 0.05 '.' 0.1 'n.s' 1

## Discussion

Here, functional developmental trajectories of the human brain were examined in the largest longitudinal perinatal fMRI dataset to date, comparing full-term and preterm populations. We examined increasing and decreasing RSFC trajectories at both global and network levels and differentiated preterm from full term cases on timing and extent of change. Studying preterm birth using two longitudinal frameworks, post-conceptional GA and birth-centered age, has the advantage of disambiguating select underlying causes of preterm delivery-related developmental differences. The centering of development at birth (birth-centered age), aligns the groups by major sensorimotor, physiological and chemical shifts that occur during the birth transition. The experiences of the newborn brain necessarily influence the expansion and pruning of neural networks. Placing extrauterine experiences on time-shifted post-conceptual maturation provides a comparison in which group brain differences are evaluated at the time the extrauterine world transition occurs, irrespective of underlying age-related programming. It also allows for isolating neural changes in the period leading up to birth, which some have suggested may differ in the preterm brain ^28^. In contrast, aligning the preterm and term-born cases by post conceptual age has the benefit of chronological synchrony, a better method for evaluating age-related programming effects. Here, we describe unique group trajectories, between our groups, within each of these frameworks.

The burst of brain development at birth in the term-born group (**Figure 1d**) is consistent with well-established biological processes underlying the perinatal transition. In late gestation, the fetus undergoes extensive physiological preparation for extrauterine life, followed by rapid adaptation to the dramatic shift in sensorimotor experience after birth. This transition encompasses a wide range of changes, from endocrine regulation and neural network maturation to macroscale structural brain reorganization ^29^. A recent study reported a substantial jump of 21.4% in cortical gyrification and an accelerated incrase of total brain surface area ^30^ around the time of term birth. This growth spurt accounts for nearly half of the total increase occuring during the fetal period (45.0%) and approximately two-thirds of the total postnatal increase during the first 5 weeks after birth (33.6%), underscoring birth as a pivotal inflection point in human brain development ^30^. Extending these structural observations to the functional domain, our findings provide novel evidence that large-scale brain functional network organization undergoes a parallel developmental inflection at birth. When cases are zero-aligned at birth, term-born infants exhibit a pronounced birth-related inflection in functional synchrony, consistent with the structural growth burst reported in prior anatomical studies. However, this burst was attenuated following correction for MRI condition differences, as MRI collection setting (in utero vs. ex utero) is fully aligned with the biological effect of birth that we aim to study. This methodological consideration is discussed further below.

In the context of preterm birth, our data do not show same degree of burst as seen in the term-born group. This pattern is also consistent with prior work on cortical gyrification, which suggests that preterm subjects resemble term-born fetuses in the early fetal period, but by term-equivalent age, preterm infants do not exhibit the discontinuity, that is, the “jump”, in gyrification seen in full-term cases ^30^. This absence of this developmental “jump” may also help explain well-documented cross-sectional findings of reduced gray matter volume ^31^ and decreased RSFC ^16^ in preterm relative to term-born neonates. The underlying neuronal mechanisms remain unclear. One hypothesis posits that cortical anatomical alterations in preterm cases is a product of neuronal loss of function and death (primary neuronal degneration) ^32^. For example, a study using human preterm infant autopsy materials and rabbit models demonstrated that neurogenesis continues during the third trimester but is suppressed by premature birth ^33^. Other hypotheses propose that cortical loss in preterm birth may reflect a failure of neuronal maturation, rather than cell death or axonal degeneration, and a third hypothesis suggests that altered growth in the thalamus may disrupt cortical development ^32^. However, there is still a lack of data to clarifying the causes of macroscale brain changes, how anatomical alterations relate to brain functional maturation, and how such brain changes affect downstream behavioural outcomes.

By comparison, when our groups are examined with alignment by post-conceptual age, a different group developmental pattern is observed. Prior to harmonization, no significant differences in either the pattern or degree of RSFC development are observed between preterm and full-term groups, with both groups following a near-linear trajectory. Following harmonization, however, preterm cases exhibit an accelerated increase in RSFC between 30 and 40 weeks of GA, whereas full-term infants maintain a largely linear developmental trajectory. We infer that this pattern in the preterm group likely represents a cumulative effect of postnatal experience, with preterm infants entering the extrauterine environment at varying gestational ages between 30 and 37 weeks, rather than an acceleration initiated by fetal programming. This observation is consistent with previous contrasting between in-utero and ex-utero preterms: preterm infants display substantially higher RSFC relative to postmenstrual age-matched fetuses^24^. At the network level, we identified similar pattern in selected systems, particulary in within-network connections in SMA, CPN, FPN, ATN,and VIC (**Figure 4**) when the data were aligned by post-conceptional age.

Except for the above mentioned within-network RSFC, other network-level differences mainly manifested as lower degree of increase in preterm cases than full-term: full-term group exhibited an overall increase in RSFC across the birth transition period, whereas preterm showed minimal change or even a decrease in RSFC in these systems over age. Overlapping findings across the birth-centerd and conception-aligned age were observed in networks including the sensotimotor network (SN), bilateral Insular Network (INS), Default Mode Network (DMN), auditory temporal network (ATN), and the large Visuomotor Integration and Coordination Network (VIC).

The insula is among the earliest developing cortical structures ^34^, and its functional segregation and role as a major hub have been observed as early as the neonatal period ^35,36^. The current study demonstrates altered developmental trajectories in preterm compared to full-term cases in how the insula communicates with both the ATN and the VIC, regions heavily involved in processing sensory information from the environment. The VIC represents another interesting work showing a convergent preterm effect. The VIC network comprises the cerebellum, subcortical regions and the inferior surface of the cerebrum, which contribute to sensory processing, including visual, auditory, and olfactory information, as well as memory-related functions. The cerebellum is the most rapidly developing brain structure during the third trimester ^37,38^, and exhibits very early functional connectivity. In our previous work, the cerebellum was identified as an emerging hub in the fetal brain ^39^. Postnatally, the cerebellum shows robust connection with subcortical regions, which are in turn extensively connected to the sensorimotor, auditory and visual cortices in neonatal brains ^35,40^, as is reflected by the large VIC network identified in the present data. Indeed, reduced cerebellar RSFC has been reported in preterm infants ^41^, and in those exposed to adverse gestational conditions, such as high maternal prenatal stress ^42^.

We also observed converging differences between preterm and full-term cases in RSFC related to DMN and other primary cortices including SN and ATN. The heightened sensibility of primary cortices to preterm-related insults is perhaps expected, given that these regions undergo rapid maturation during the perinatal period. However, the DMN, a higher-order association network important for self-referential thought ^43^, exhibits widespread differences in RSFC, both within the network and in its connectivity with other networks, between the preterm and full-term groups. These differences suggest that DMN maturation may begin earlier than previously anticipated, though the underlying mechanisms remain to be elucidated. It is possible that these areas represent alternative developmental pathways, or that they emerge as a response to early postnatal experience. Further studies will be needed to disentangle why selected systems show alterations in preterm group, and to clarify the functional significance of these regional effects.

Several important caveats are worth highlighting. Most notably, there are inherent limitations in the datasets, including the lower signal-to-noise ratio in fetal compared with infant fMRI scans, and heterogeneity in data acquisition and preprocessing strategies across cohorts and between *in utero* and *ex utero* conditions. These challenges are further compounded when data are aligned to the birth event, as the covariate we aim to control (differences in MRI acquisition condition), becomes collinear with the variable of primary interest (the birth event itself). Unfortunately, no ground truth exists to isolate the effect of imaging condition, as there is no constant reference variable against which its influence can be assessed. To account for this, we implemented 2 strategies in our analysis to mitigate the impact of different temporal singal-to-noise ratio between datasets: (1) In our RSFC analysis, we used the z-values of RSFC for all subsequent analyses to correct for the variance differences. (2) We applied a harmonization approach to eliminate the effects of this variance difference before conducting our statistical analyses. To maximize transparency, we have included both pre- and post-harmonization results throughout the manuscript and Supplementary Materials, allowing readers to directly evaluate the data and draw their own informed interpretations. Furthermore, we applied strict exclusion criteria based on image quality (see Methods) and conducted a sensitivity analysis using an image-quality– matched subset to replicate our main results (see **Supplementary Materials Figure S1**). Another consideration is that this study invites many opportunities for examining individual circumstances of the pregnancy or postpartum environment that may alter individual trajectories. We established that mode of delivery and biological sex were not important determinants in this sample, however, we did not address many putative infunces, such as maternal health conditions or time spent in the NICU, both of which influence brain and behavioral development ^44,45^.

The present findings represent an encouraging first step toward a deeper understanding of brain prematurity and of “birth” as a transformative event. As ongoing and future efforts yield increasing amounts of high-quality perinatal MRI data, we anticipate that improved brain charts for the perinatal period will be iteratively refined using multimodal neuroimaging phenotypes. Follow-up studies should assess alterations in RSFC related to the targeted regions and neural pathways. Such robust models may improve understanding of neural effects underlying behavioral outcomes associated with prematurity. It is also possible that these data may be used to identify sensitive periods and/or modifiable targets for intervention.

In summary, leveraging a unique perinatal neuroimaging dataset, our analyses reveal differentiable developmental trajectories in large-scale functional network synchrony between full-term and preterm cases. When data are aligned at birth, preterm infants exhibit generally lower RSFC than their full-term counterparts. In contrast, when data are aligned by post-conceptional age, preterm infants display an accelerated increase in RSFC beginning around 30 gestational weeks. These trajectory differences are further reflected in selected functional networks, suggesting that distinct developmental strategies may operate across different neural systems in the context of prematurity. Notably, the mode of delivery and sex showed minimal influence on these trajectories. Together, these findings provide unprecedented insight into how prematurity and early exposure to the extrauterine environment reshape the pace and sequence of network maturation, advancing our understanding of both neurotypical and preterm brain development.

## Methods

### Participants and Datasets

#### *Dataset 1:* Developing Human Connectome Project (dHCP)

Data used in this study were obtained from the fourth data release of the dHCP (http://www.developingconnectome.org/project/). The study was approved by the UK Health Research Authority (Research Ethics Committee reference number: 14/LO/1169) and parental written consent was obtained. Recruitment took place at the Evelina Newborn Imaging Centre, St Thomas’ Hospital in London, UK. Main inclusion criteria were 1) pregnant woman with fetal age estimated from the last menstrual period between 22 and 44 weeks and 2) living infants between 23 and 44 weeks of GA. Main exclusion criteria were 1) contraindication to undergo MRI scanning in the mother of the infant, 2) expert medical advice against MRI due to health conditions in PT infants and 3) failed communication regarding study procedures and consent. A total of 263 fetal fMRI scans and 811 infant fMRI scans were successfully downloaded. Pregnancies with multiple fetuses (i.e., non-singleton pregnancies) were excluded from this study.

#### *Dataset 2:* Perinatal Imaging of Neural Connectivity (PINC) project

This study included 157 fetal fMRI scans, and 57 infant fMRI scans collected between 2013 and 2018, as part of the Perinatal Imaging of Neural Connectivity (PINC) project at Wayne State University (WSU). The WSU Institutional Review Board approved all study procedures, and informed written consent was provided by participating pregnant people. Eligibility criteria included singleton pregnancy, maternal age 18–40, no suspected central nervous system abnormality as determined by 20-week ultrasound, and no contraindication for MRI. Select cases were scanned 1-3x longitudinally, between post-conceptual 25 weeks and 55 weeks of GA. All included scans underwent preliminary quality control, ensuring (1) a minimum of 105 volumes and (2) complete brain coverage without significant artifacts or distortions.

### FMRI acquisition

#### Dataset1: dHCP

##### Fetal fMRI

Following the acquisition protocol described in reference^46^, dHCP fetal fMRI data were acquired with a Philips Achieva 3T system (Best, NL) using a 32-channel cardiac coil at St Thomas’ Hospital London. Each dataset included 350 volumes (48 slices per volume) using a single-shot echo EPI sequence (TR/TE = 2200/60 ms), with a 144 × 144 matrix, 2.2 mm isotropic resolution, MB factor = 3, and SENSE factor = 1.4. Datasets with poor image quality were excluded from the public release.

##### Infant fMRI

Following the acquisition protocol reported in reference ^47,48^, dHCP infant fMRI data were collected on a 3T Philips Achieva system equipped with a dedicated neonatal imaging setup located within the neonatal intensive care unit at Evelina London Children’s Hospital. A high–temporal-resolution multiband EPI sequence optimized for neonates (TE = 38 ms; TR = 392 ms; 2300 volumes; MB factor = 9; 2.15 mm isotropic) was acquired over a 15-minute scan. No in-plane acceleration or partial Fourier was applied.

#### Dataset2: PINC

##### Fetal fMRI

As described in our previous publication^49^, PINC fetal fMRI was acquired using a 3T Siemens Verio 70 cm open-bore system with an abdominal 4-channel Siemens Flex coil. FMRI data were acquired using one of two protocols: (i) 12-min single-echo fMRI: TR/TE = 2000/30 ms; resolution = 3.4 × 3.4 × 4 mm^3^, flip angle: 80 degrees. (ii) 12-min multi-echo (ME) fMRI: TR = 2000ms; TE = 18, 34, 50ms (3 echoes); flip angle: 83 degrees; voxel size: 3.5 x 3.5 x 3.5 mm^3^. Scans were repeated when time permitted.

##### Infant fMRI

As previous described ^49^, PINC infants were scanned on the same 3T Siemens Verio 70 cm open-bore system with a 32-channel head coil, using one of three acquisition protocols: (i) 12-min ME-fMRI: TR = 2000 ms; TE = 13, 26, 39 ms; flip-angle: 83 degrees; slice-gap: none; voxel-size: 3.5 × 3.5 × 3.5 mm^3^; matrix-size: 64 × 64 × 39 voxels. (ii) 12-min multi-band (MB) ME-fMRI scan: TR = 1500 ms; TE = 15, 31, 46 ms; flip-angle: 83 degrees; slice-gap: none; voxel-size: 2.9 × 2.9 × 2.9 mm^3^; matrix-size: 64 × 64 × 48 voxels, multi-band factor = 2. (iii) 7-min MB ME fMRI scan: TR = 1000 ms; TE = 14.6, 36.68, 58.76 ms; flip-angle: 52; slice-gap: none; voxel-size: 2.5 × 2.5 × 2.5 mm3; matrix-size: 80 × 80 × 44 voxels, multi-band factor = 4.

### FMRI preprocessing

#### Dataset1: dHCP

FMRI data preprocessed by the dHCP scientific collaborators were used for the present work.

##### Fetal fMRI

For complete information on fetal fMRI preprocessing, we refer to the dHCP publications ^46^. Briefly, the preprocessing pipeline included: (1) MB-SENSE image reconstruction, to generate high-quality images from the multi-band acquisition while correcting for coil sensitivity profiles; (2) dynamic distortion correction, using the phase information from the reconstructed gradient echo EPI images acquired for fMRI; (3) motion correction, including slice-to-volume motion correction ^50^; (4) native-to-template registration comprising native functional-to-structural alignment, structural-to-age-matched template mapping, and stepwise template-to-template registration to the 36-week GA template; and (5) temporal filtering, involving regression of multiple artifact-related nuisance regressors. The data we downloaded were preprocessed in native space, along with the corresponding warp mappings to the 36-week GA template. We applied these warps to the data and subsequently downsampled the images from 0.5 mm isotropic resolution to 1.5 mm isotropic.

##### Infant fMRI

For details on infant fMRI preprocessing, please see reference^47^. The main parts of the pipeline include: (1) field map preprocessing, to estimate the susceptibility distortion field and align it with the functional data; (2) susceptibility and motion correction, including slice-to-volume motion correction, dynamic susceptibility distortion correction, and estimation of motion-related nuisance regressors; and (3) denoising, involving estimation of artefact-related nuisance regressors and regression of all nuisance signals from the functional data. We then normalized the preprocessed data to the dHCP 36-week GA template (1.5 mm isotropic resolution) using a two-step registration process: functional-to-neonatal template alignment followed by neonatal template-to-36-week template mapping, both estimated with Advanced Normalization Tools (ANTs v2.4.3)^51^.

#### Dataset2: PINC

##### Fetal fMRI

Details of the fetal preprocessing pipeline have been reported previously^49^. the pipeline included: (1) automatic fetal brain segmentation using a deep learning model^52^, guided by a manually drawn reference mask and refined into a precise, individualized 4D mask through iterative transformation and masking steps; (2) motion parameter estimation and three-step frame censoring based on Sørensen–Dice coefficient, framewise displacement, and DVARS thresholds; (3) optimal combination across echoes using T2*-weighted averaging^53^; (4) denoising using individual-level ICA implemented in FSL MELODIC^54^, (5) manual reorientation in SPM ^55^, and (6) normalization to the 36-week GA template (isotropic 1.5mm), ensuring consistency with the dHCP cohort.

##### Infant fMRI

Infant brain preprocessing followed prior published procedures ^49,56^, beginning with FSL’s Brain Extraction Tool (BET, ^57^) and motion correction using “FSL mcflirt”. Volumes with a root-mean-square DVARS value greater than 50 were identified as outliers and censored, along with one preceding and two subsequent frames. Motion-corrected data were subsequently combined across echoes, denoised using TE-Dependent Analysis (Tedana v22.0.1, ^58,59^), normalized to the dHCP 36-week GA template (isotropic 1.5mm).

All normalized data in standard space, including both fetal and infant scans from the dHCP and PINC datasets, were smoothed using a 3 mm Full Width at Half Maximum (FWHM) kernel using *fslmaths*. A further step of denoising, CompCor denoising, were performed on all smoothed data using Functional Connectivity Toolbox (CONN21.a) ^27,60^. Specifically, during CompCor, the regression of potential confounding effects includes white matter timeseries (5 CompCor noise components), CSF timeseries (5 CompCor noise components), and linear trends (2 factors) within each functional run, followed by bandpass frequency filtering of the BOLD timeseries between 0.008 Hz and 0.09 Hz to remove low frequency temporal drifts and high frequency physiological and scanner-related artifacts ^61^.

### FMRI quality assurance

The root-mean-square of voxel-wise differentiated signal (DVARS) was calculated for all preprocessed and normalized scans. DVARS distribution for each dataset are provided and **Figure 6**. As shown in **Figure 6**, data quality was comparable across the PINC and dHCP cohorts for both fetal and infant scans. As expected, fetal scans exhibited overall higher DVARS values than infant scans, reflecting greater motion during fetal imaging.

**Figure 6.**
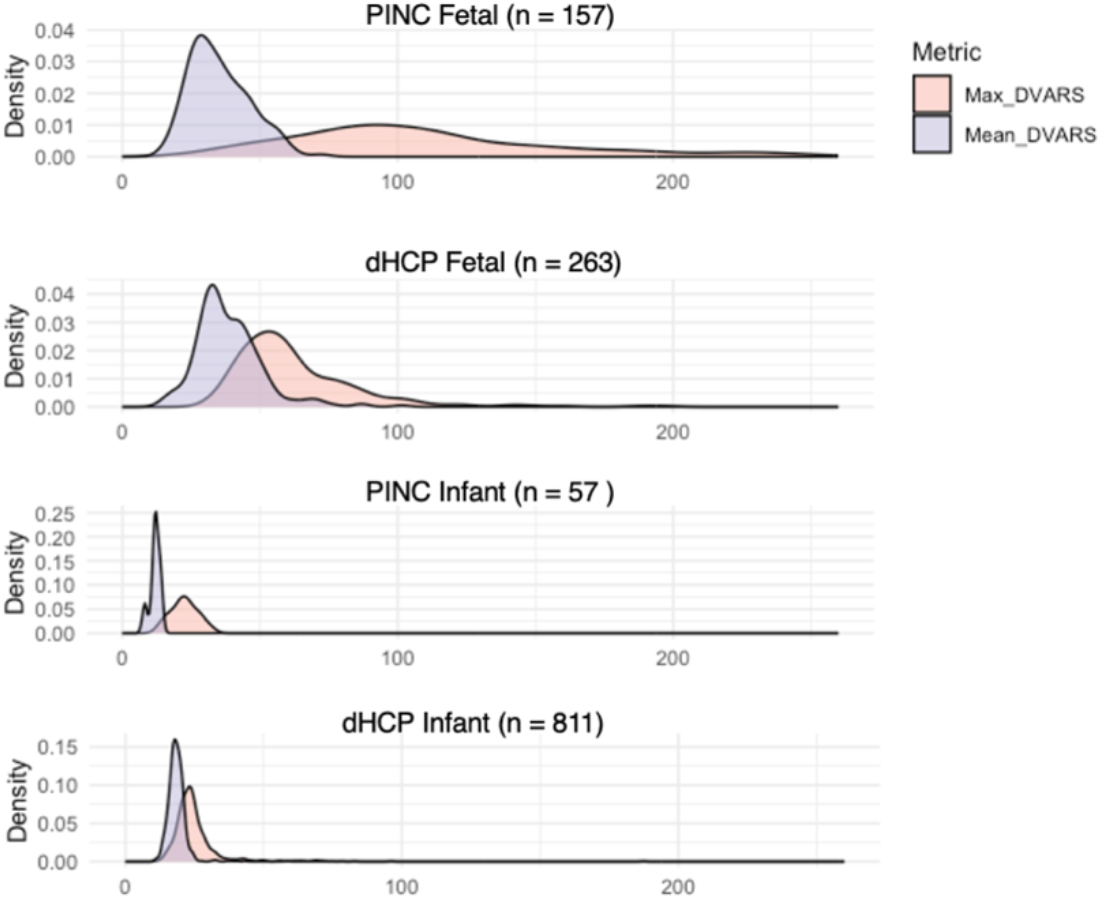
Density distributions of DVARS across fetal and infant datasets before quality exclusion. Density plots show the distribution of maximum DVARS (pink) and mean DVARS (purple) for four datasets, displayed from the top to bottom: PINC fetal, dHCP fetal, PINC infant, and dHCP infant.

MRI data were further excluded based on DVARS thresholds (mean DVARS < 50 and maximum DVARS < 150). Following this quality control, the final dataset comprised 174 scans from 143 unique subjects in PINC and 857 scans from 727 unique subjects in dHCP, for a total of 1,031 scans (see GA at scab across cohorts at Supplmentary **Figure S4**).

### Functional connectivity matrix construction

A subset of 200 fetal and infant datasets (50 from each dHCP and PINC fetal/infant cohort with lowest DVARS) was used to generate a data-driven, group-balanced functional atlas. Gray-matter-masked data were submitted to the SLIC toolbox ^62,63^ to produce 200 functional parcels. Pairwise zero-lag correlations between ROI time courses were Fisher-z transformed to generate a 200 × 200 functional connectivity matrix for each scan in CONN 21.a. Linear regression models were used to test age effects for RSFC in CONN second-level analysis, controlling for the group-related variance in age. Regression analyses were thresholded at p<0.05, FDR corrected. Average RSFC that exhibited significant positive or negative associations with age (RSFC connection threshold p < 0.05) were then extracted to derive growth plots.

### Harmonization on MRI condition effects

To account for non-biological variance introduced by different MRI conditions (i.e., *in-utero* vs. *ex-utero* effects) and datasets (dCHP vs. PINC), we applied a longitudinal harmonization technique to the our RSFC measures using the R package LongComBat ^26^. This tool has demonstrated superior efficacy in detecting longitudinal changes compared to an alternative harmonization method, cross-sectional ComBat, while also providing better control of type I error rates than unharmonized data that incorporates scanner as a covariate ^26^.

### Delivery records

#### Dataset1: dHCP

Delivery mode data were recoded into a binary variable representing vaginal or caesarean delivery. Instrumental deliveries (forceps or ventouse) were classified as vaginal, while semi-elective caesarean sections were classified as caesarean.

#### Dataset2: PINC

Delivery mode data for PINC were extracted from medical records.

### Statistical analysis

Growth trajectories were modeled using generalized additive mixed-effect models (GAMMs) in R with the “*gamm4*” function, applied to global average RSFC as well as within- and between-network RSFC pairs. Models followed the specification: RSFC ∼ 1 + s(GA) + (1|Subject), and were fit separately for full-term and preterm groups. The first derivative of each fitted curve was then calculated to quantify the rate of change in RSFC over time.

To understand the development patterns for each network, we also conducted GAMM analyses on each within- and between-network connectivity across 8 global networks (N= 36 possible associations), using the same approach as described above. The network-level RSFC was calculated by averaging the ROI-by-ROI RSFC values that showed significant positive or negative associations with age.

## Acknowledgement

We are grateful to the families who generously supported these projects.

## Data availability

This paper analyzes existing available data from the developing Human connectome Project (dHCP) and Perinatal Imaging of Neural Connectivity (PINC) project. The dHCP is an open-access project. The imaging and collateral data used in this study were included in the 4th dHCP data release, which can be downloaded by registering at https://data.developingconnectome.org/. PINC data used in this study will be made available via https://ndar.nih.gov/ and/or accessed upon direct request to M.E. Thomason. The codes for PINC fetal MRI preprocessing is available on Zenodo: 10.5281/zenodo.13905953. 10.5281/zenodo.13984128.

