## Supplementary Materials for "Full-Term and Preterm Differences in Human Brain Functional Connectivity Development Across Birth"

**Table S1. Sensitivity Analysis: Group compassions across demographic factors, birth outcomes, and MRI data quality.**

| Cohort Characteristics | PRETERM (N = 244) | | FULL TERM (N = 787) | | p-value |
|---|---|---|---|---|---|
| | Mean (SD) | Range | Mean (SD) | Range | |
| Fetal sex, N (%) | | | | | 0.649 |
| F | 106 (43.4%) | | 112 (45.9%) | | |
| M | 138 (56.6%) | | 132 (54.1%) | | |
| Post-conceptional GA (weeks) at scan | 36.65 (4.89) | (24.14, 51.57) | 40.19 (5.48) | (21.86, 54.57) | <0.001 |
| Post-conceptional GA (weeks) at birth | 32.33 (3.57) | (23.00, 37.00) | 39.60 (1.22) | (37.14, 42.14) | <0.001 |
| Birth-centered age (weeks) | 4.32 (5.85) | (-12.57, 20.57) | 0.59 (5.39) | (-17.85, 13.86) | <0.001 |
| Scan Type | | | | | 0.280 |
| Fetal | 27 (11.1%) | | 36 (14.8%) | | |
| Neonatal | 217 (88.9%) | | 208 (85.2%) | | |
| Delivery method | | | | | <0.001 |
| Vaginal | 113 (46.3%) | | 168 (68.9%) | | |
| C-section | 129 (52.9%) | | 71 (29.1%) | | |
| Missing | 2 (0.8%) | | 5 (2.0%) | | |
| Fetal MRI Characteristics | | | | | |
| Mean DVARS | 19.28 (7.15) | (10.89, 49.34) | 19.38 (7.44) | (10.15, 49.89) | 0.881 |
| Max DVARS | 29.61 (24.48) | (11.38, 142.99) | 30.97 (20.81) | (15.35, 144.36) | 0.510 |

#### Figure S1. Sensitivity analysis

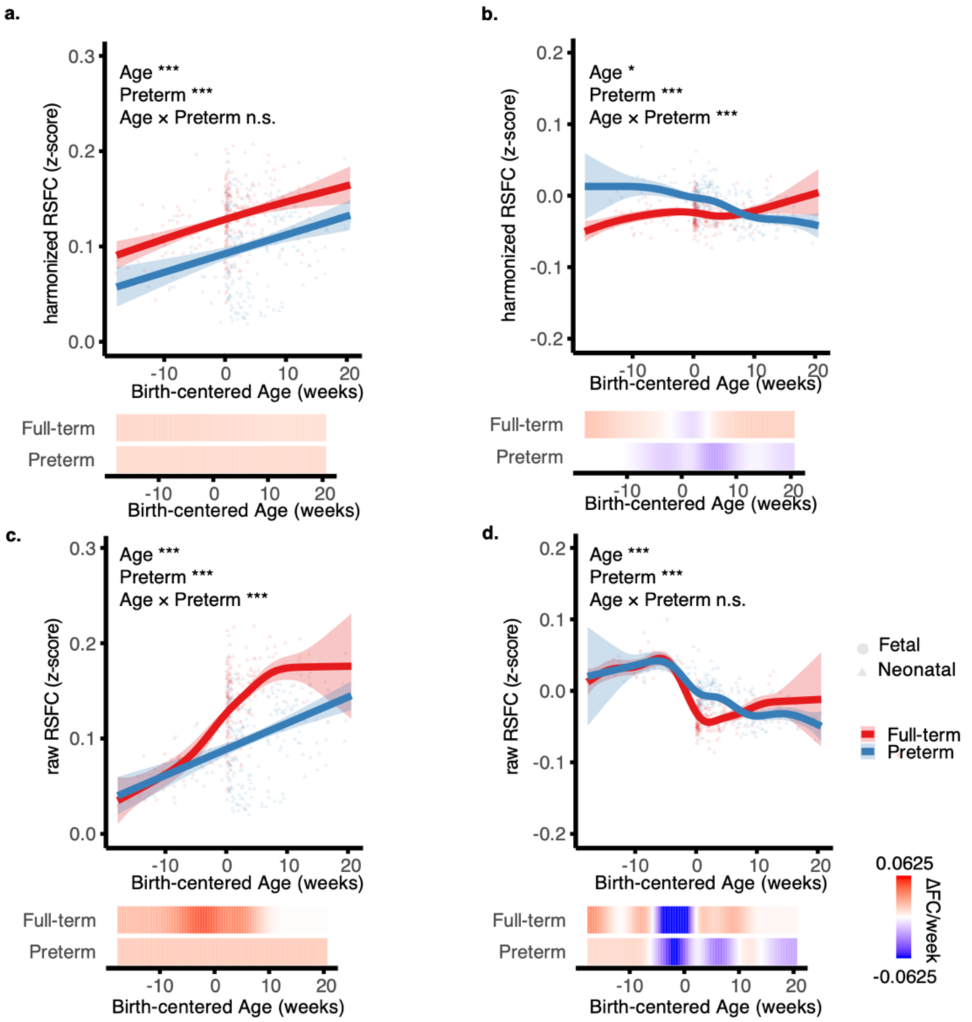

### Figure S2. Pre-harmonization plots with birth-centered age

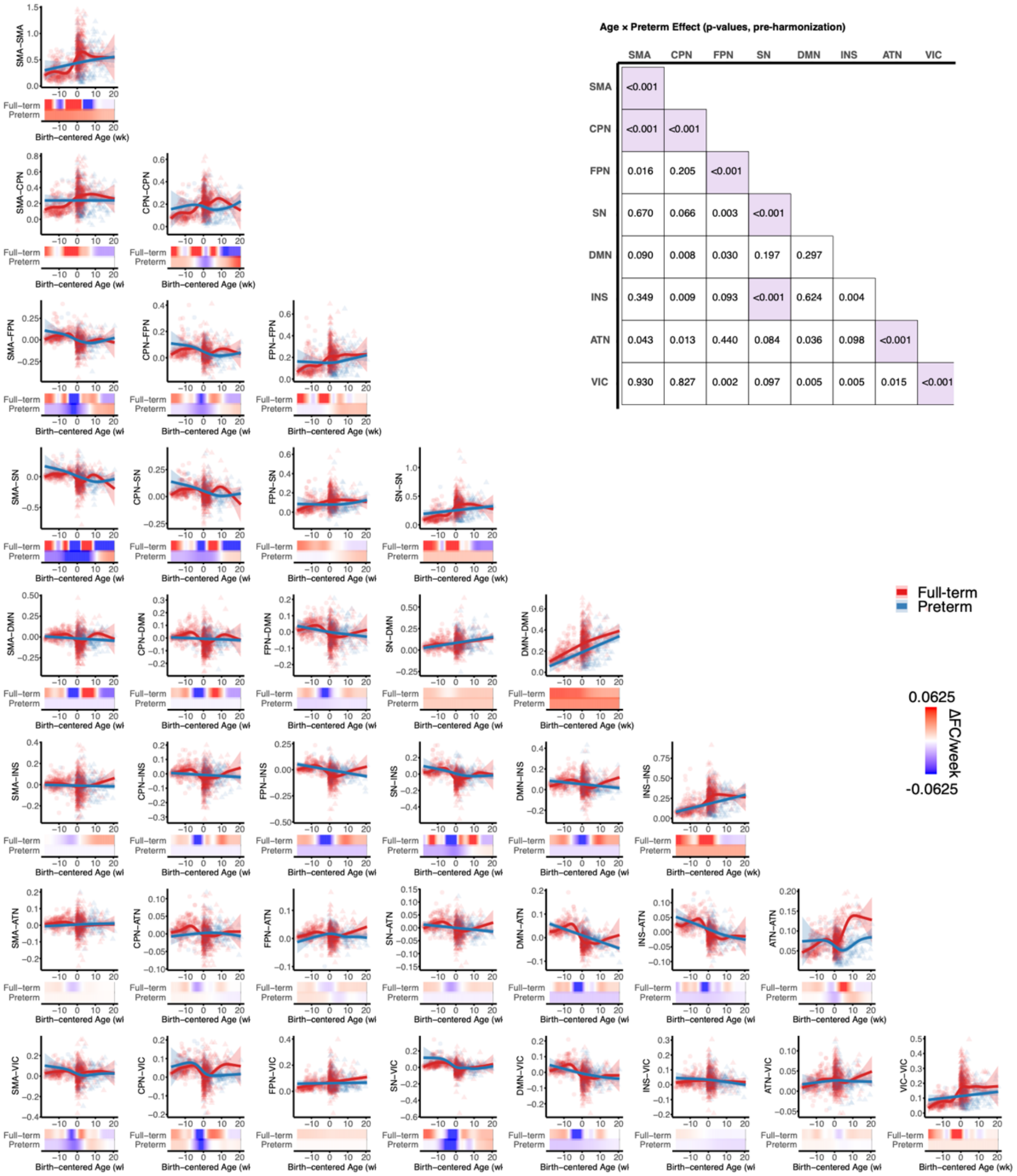


**Age × Preterm Effect (p-values, pre-harmonization)**

| | SMA | CPN | FPN | SN | DMN | INS | ATN | VIC |
|---|---|---|---|---|---|---|---|---|
| SMA | <0.001 | | | | | | | |
| CPN | <0.001 | <0.001 | | | | | | |
| FPN | 0.016 | 0.205 | <0.001 | | | | | |
| SN | 0.670 | 0.066 | 0.003 | <0.001 | | | | |
| DMN | 0.090 | 0.008 | 0.030 | 0.197 | 0.297 | | | |
| INS | 0.349 | 0.009 | 0.093 | <0.001 | 0.624 | 0.004 | | |
| ATN | 0.043 | 0.013 | 0.440 | 0.084 | 0.036 | 0.098 | <0.001 | |
| VIC | 0.930 | 0.827 | 0.002 | 0.097 | 0.005 | 0.005 | 0.015 | <0.001 |

**Figure S3. Pre-harmonization plots with post-conceptional age**

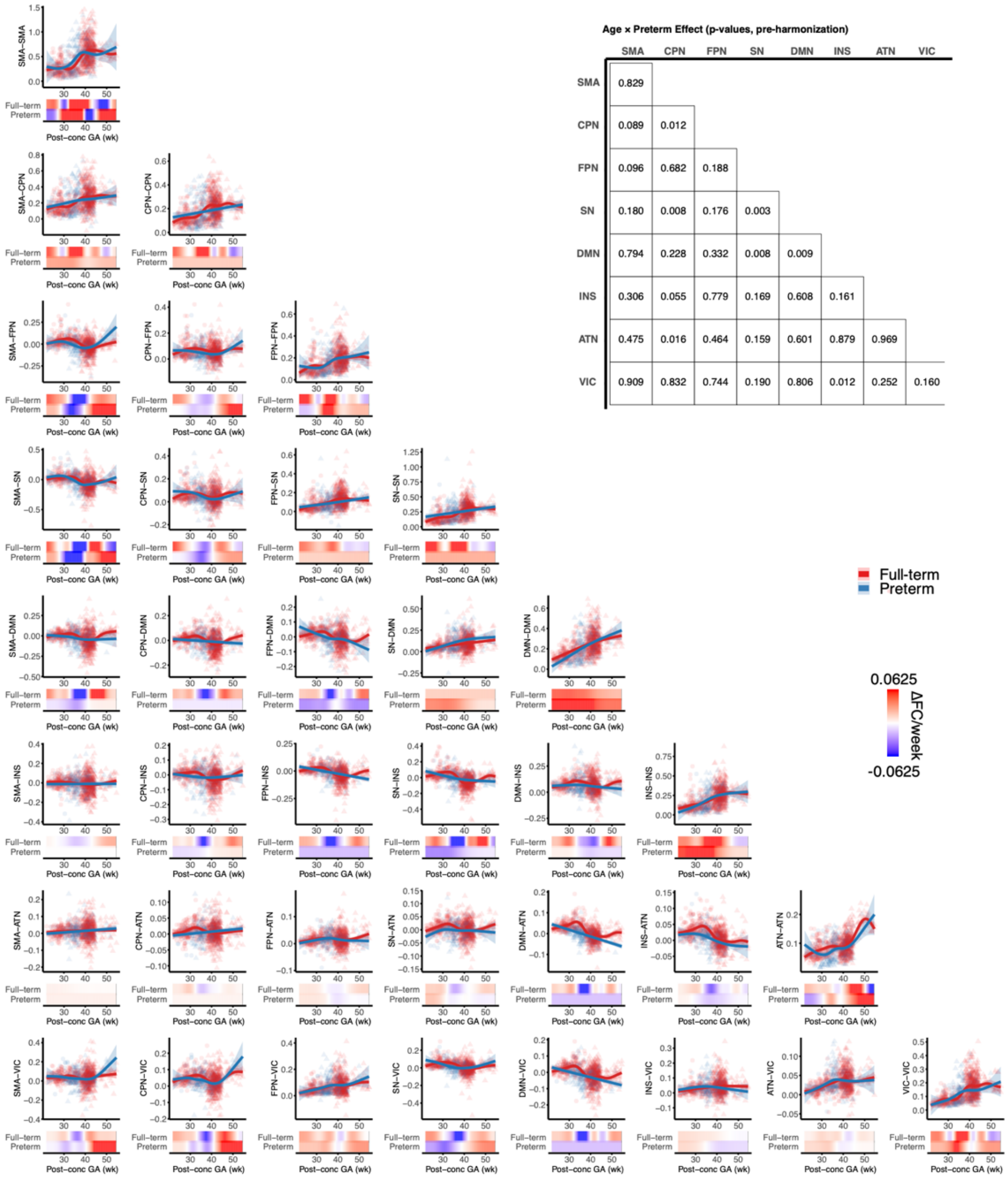


**Age × Preterm Effect (p-values, pre-harmonization)**

| | SMA | CPN | FPN | SN | DMN | INS | ATN | VIC |
|---|---|---|---|---|---|---|---|---|
| SMA | 0.829 | | | | | | | |
| CPN | 0.089 | 0.012 | | | | | | |
| FPN | 0.096 | 0.682 | 0.188 | | | | | |
| SN | 0.180 | 0.008 | 0.176 | 0.003 | | | | |
| DMN | 0.794 | 0.228 | 0.332 | 0.008 | 0.009 | | | |
| INS | 0.306 | 0.055 | 0.779 | 0.169 | 0.608 | 0.161 | | |
| ATN | 0.475 | 0.016 | 0.464 | 0.159 | 0.601 | 0.879 | 0.969 | |
| VIC | 0.909 | 0.832 | 0.744 | 0.190 | 0.806 | 0.012 | 0.252 | 0.160 |

**Figure S4. GA at MRI scan across PINC and dHCP cohorts. (A)** Distribution of post-conceptual GA at scan for the PINC (pink) and dHCP (purple) datasets. Vertical dashed lines mark the cohort mean GA at scan. **(B-C)** Lollipop plots depicting individual participants from the PINC (B) and dHCP (C) cohorts. Each line corresponds one participant, with points representing scan times. Fetal scans are shown in dark grey and infant scans in light grey. Pink dots indicate GA at birth for participants.

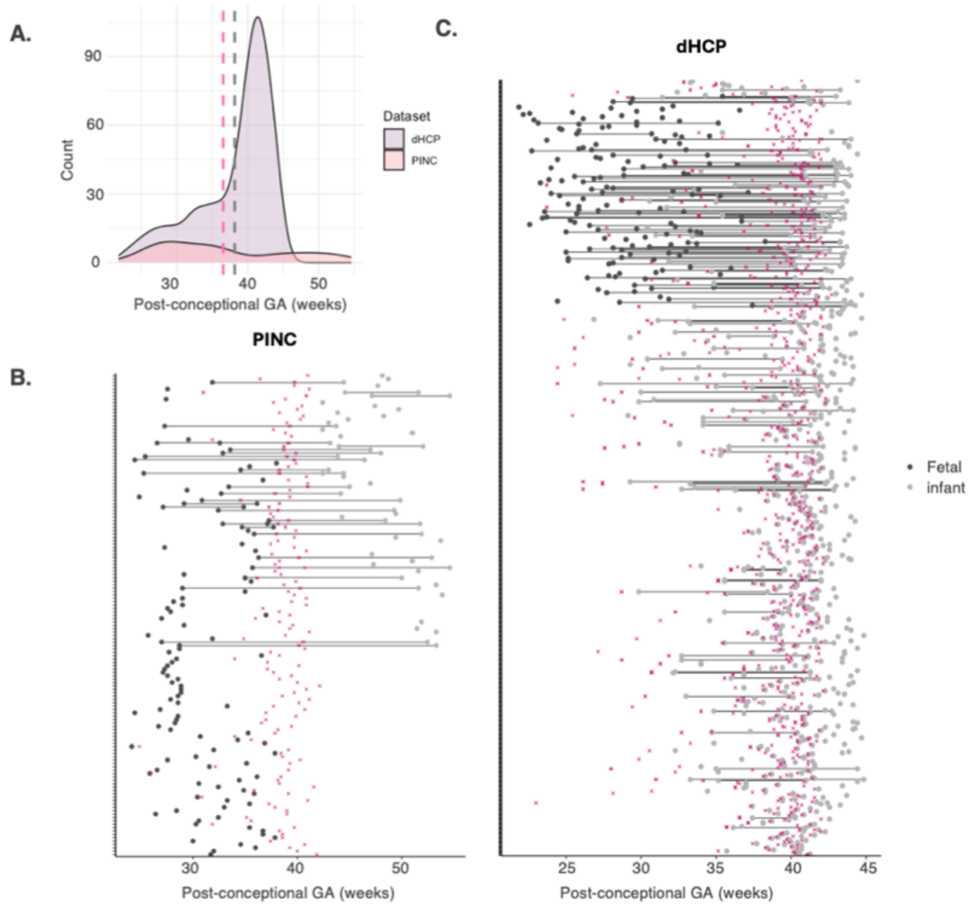
